# Genetic modification of the ant *Lasius niger* using CRISPR-Cas9 technology

**DOI:** 10.1101/2022.05.02.490261

**Authors:** Mauno Konu, Jonna Kulmuni, Lumi Viljakainen

## Abstract

CRISPR-Cas9 has become one of the most prominent gene editing tools available and it has been utilized in various organisms from bacteria to fungi, plants, and animals. In this study, we developed a CRISPR-Cas9 protocol for the black garden ant *Lasius niger*, a common and easily available study species for lab and field environments. To create indel mutations using CRISPR-Cas9 in *L. niger*, we targeted three different locations in a well-studied eye pigmentation gene *cinnabar*, generating several mutations that disrupt the ommochrome biosynthesis pathway and result in the lack of this pigment and therefore, abnormal eye coloration in adult workers. We also developed a protocol to collect *L. niger* eggs, inject them with CRISPR-Cas9 construct, and rear the eggs into mature adult workers with assistance of nursing workers. We demonstrated for the first time with *L. niger* that CRISPR-Cas9 is an excellent tool to create targeted mutations for this species. Our protocol can be referred to when developing similar studies for other species of ants and eusocial insects.

## INTRODUCTION

Gene modification can be used to study the function of genes by disrupting the gene sequence and observing changes caused in the phenotype or fitness. Alternatively, it can be used to add new genes to the genome and thus new functions or repair nonfunctional genes such as hereditary diseases (Sander & Joung, 2014). CRISPR-Cas9 is the latest addition to the gene modification toolkit that has gained popularity due its simplicity and cheap price (Cui, Sun & Yu, 2017).

CRISPR-Cas9 technique has been widely used from bacteria to plants and animals. In insects this technique has been successfully used in different model species. The first was the fruit fly *Drosophila melanogaster* in 2013 (Gratz *et. al*., 2013) and after that, for example the silk moth *Bombyx mori* and the honeybee *Apis mellifera* (Cui, Sun & Yu, 2017). CRISPR-Cas9 has been utilized in at least three ant species prior to this study: twice in *Harpegnathos saltator* (Yan *et. al*., 2017; Sieber *et. al*., 2020) and once in *Ooceraea biroi* (Trible *et. al*., 2017) and *Solenopsis invicta* (Chiu *et. al*., 2020).

All these studies noted that the distinctive behavior of ants such as offspring’s helplessness and need for care, interaction and communication between individuals and aggressiveness towards strangers cause difficulties to rear genetically modified eggs into adults. Eggs and larvae require workers’ attention and e.g. individuals with modified olfactory receptors had lower fitness in the colony (Trible *et. al*., 2017). In these studies, different methods were developed to rear larvae into adults (Trible *et. al*., 2017; Yan *et. al*., 2017; Sieber *et. al*., 2020) or success of the gene modification was detected by sequencing at the larval stage (Chiu *et. al*., 2020).

This study aimed to develop a CRISPR-Cas9 protocol for *Lasius niger*, the black garden ant. *L. niger* is a widely distributed ant species ranging all of Europe and temperate Asia (Haatanen, Ooik & Sorvari, 2015). It is the most common ant species in European urban areas. Like all other ant species, it is eusocial (Ward, 2014) with colonies consisting of cooperating individuals: up to 10 000 sterile workers and one reproductive queen that together take care of the brood laid by the queen (Haatanen, Ooik & Sorvari, 2015). The queens have the longest recorded lifespan of all eusocial insects, as much as 28 years while workers live from one to two years (Kramer, Schaible & Scheuerlein, 2016). Black garden ant is commonly used as a laboratory study species because queens are easy to collect during mating flights, artificial colonies are easy to establish and maintain in laboratory conditions and the species is not very aggressive. A colony survives at room temperature and is not picky about food sources if carbohydrates and proteins are available. During winter the species overwinters, which is easy to arrange at refrigerator temperature.

We chose to modify the *cinnabar* gene that plays a role in the formation of insect eye color. Most of the wide spectrum of insect color is caused by three pigment groups: melanins, ommochromes and pteridines (Brent & Hull, 2019; Okude & Futahashi, 2021). Melanins produce black and brown colors, ommochromes brown and red colors and pteridines red, yellow and blue colors. The colors and patterns found in insects are produced by mixing and regulating the relative amounts of these pigments. Even though ommochromes produce brown color, the genes are named after the shades of red. Mutations interrupt the production of brown color in different stages of the biosynthesis and only red pigments from uninterrupted pteridine biosynthesis pathway can accumulate into the pigment granules. One of these genes, *cinnabar*, is heavily studied, and it has been modified with CRISPR technology in different species such as the parasitoid wasp *Nasonia vitripennis* (Li *et. al*., 2017) and the planthopper *Nilaparvata lugens* (Xue *et. al*., 2018), where dysfunctional *cinnabar* results in red eyes. *cinnabar* and other genes affecting eye coloration pigments are excellent targets for studies in which genes of a new species are modified for the first time, because mutations are not lethal and they are easy to observe by eye (Xue *et. al*., 2018).

Here, we report a successful gene modification protocol from collecting the ants from the wild to verification of mutants in the laboratory. We also give details on the rate of successful colony foundation in the laboratory, optimized rearing conditions for CRISPR-Cas9 injected eggs, mortality rate during the experimental procedure and success rate of the genetic modification when scored based on either phenotype or genotype. By this study, *L. niger* is the fourth ant species in which CRISPR-Cas9 technique has been successfully implemented. Methods used in this study can be utilized in future studies with *L. niger* and very likely with other ant species and eusocial insects.

## RESULTS

### Half of the collected queens managed to establish a laboratory colony

A total of 286 queens were collected to establish laboratory colonies. The first eggs emerged on the first day. The first larvae were observed 18 days after queen collection and the first pupa 25 days after collection. The first pupa hatched 32 days after collection and were at this stage still all white and immobile. The first worker matured enough to start walking 42 days after collection, after which the colony was deemed established and it was transferred from a tube to a Petri dish nest (see set up in the Material and Methods). The busiest time was 42-52 days after collection when, within ten days, 137 colonies were established. After this the emergence rate of new workers in the tubes started to decline and the last colonies were established 87 days after collection. Monitoring of the tubes was stopped 115 days after collection. Overall, 160 queens (56%) established a colony (Figure 1). In addition, 66 older colonies established in 2019 were used in this experiment.

**Figure 1:**
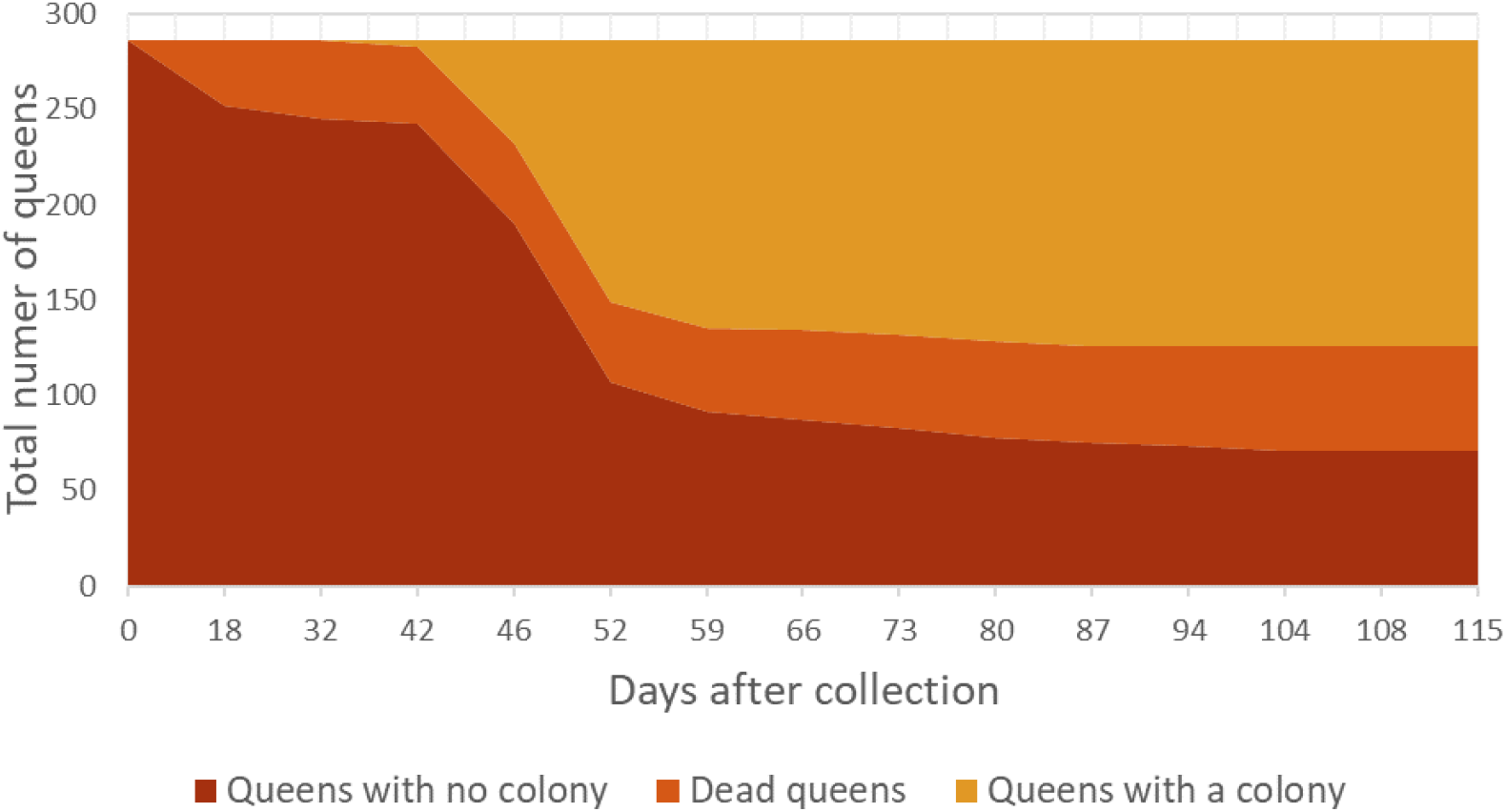
Half of the collected queens established a laboratory colony. In total 56% of all queens managed to establish a colony, 19% of the queens died during the first 115 days and 25% of the queens did not produce adult workers and therefore failed to establish a colony.

### The queens’ egg-laying rate increased towards the end of the day

We injected the CRISPR-Cas9 construct into freshly laid eggs. We performed the CRISPR-Cas9 experiment in two batches. During the first batch it was observed that the queens’ egg-laying rate increased during the day. In the second batch the egg-laying time was extended from a total of four hours to five hours. The last hours (4^th^ and 5^th^ hour) produced the most eggs (Figure 2). Likewise, in both batches the daily egg-laying rate increased during the first week and then started to decrease (Figure 3). In the span of 36 collection days in both batches (2 × 18 days, both in 24 days’ time period), a total 1636 eggs were laid by a total of 203 queens. The second batch produced more than twice as many eggs as the first batch due to the one hour longer egg-laying time. The number of eggs laid by a single queen varied a lot between individuals (Figure 4).

**Figure 2:**
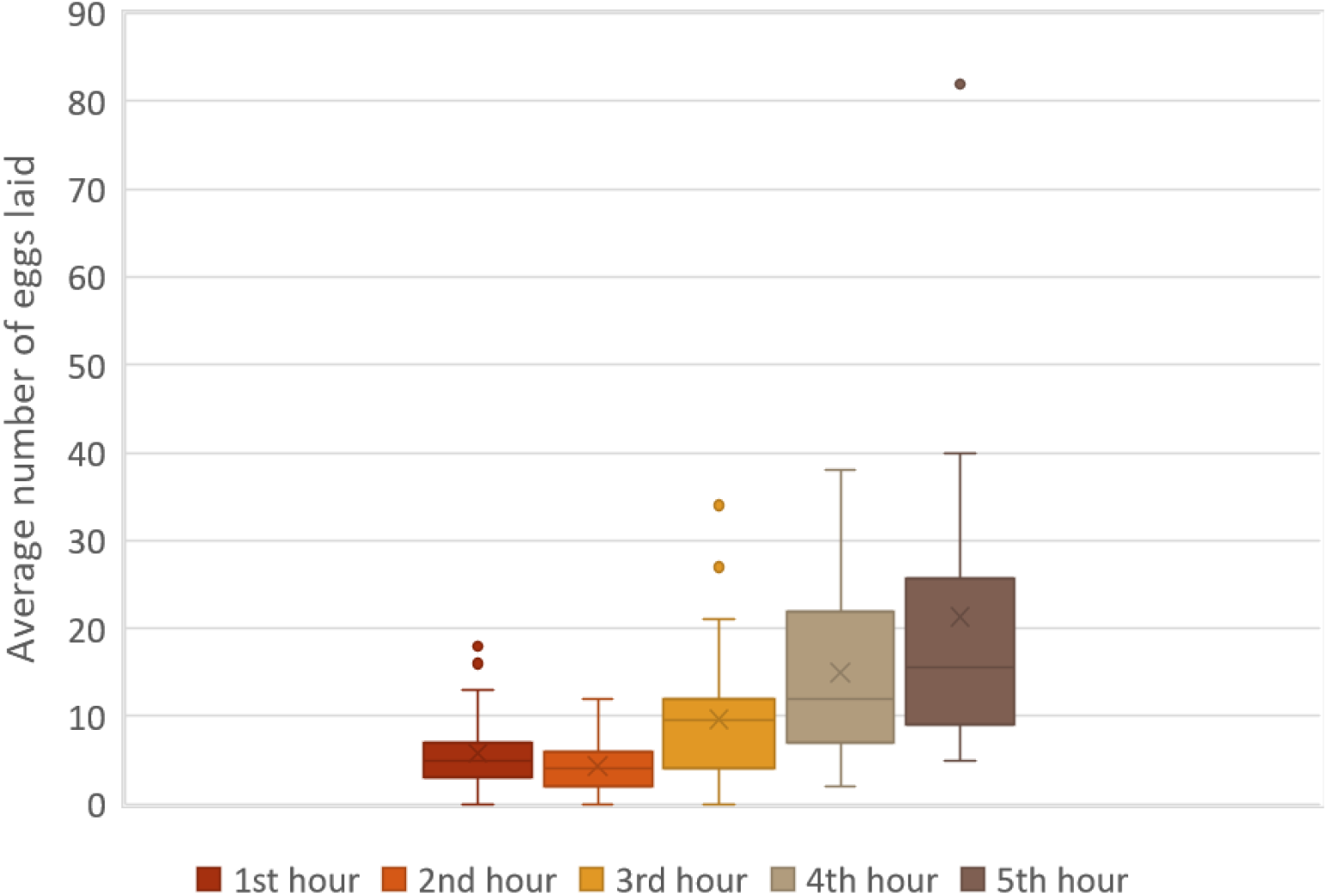
The average number of eggs laid increased towards the end of the day.

**Figure 3:**
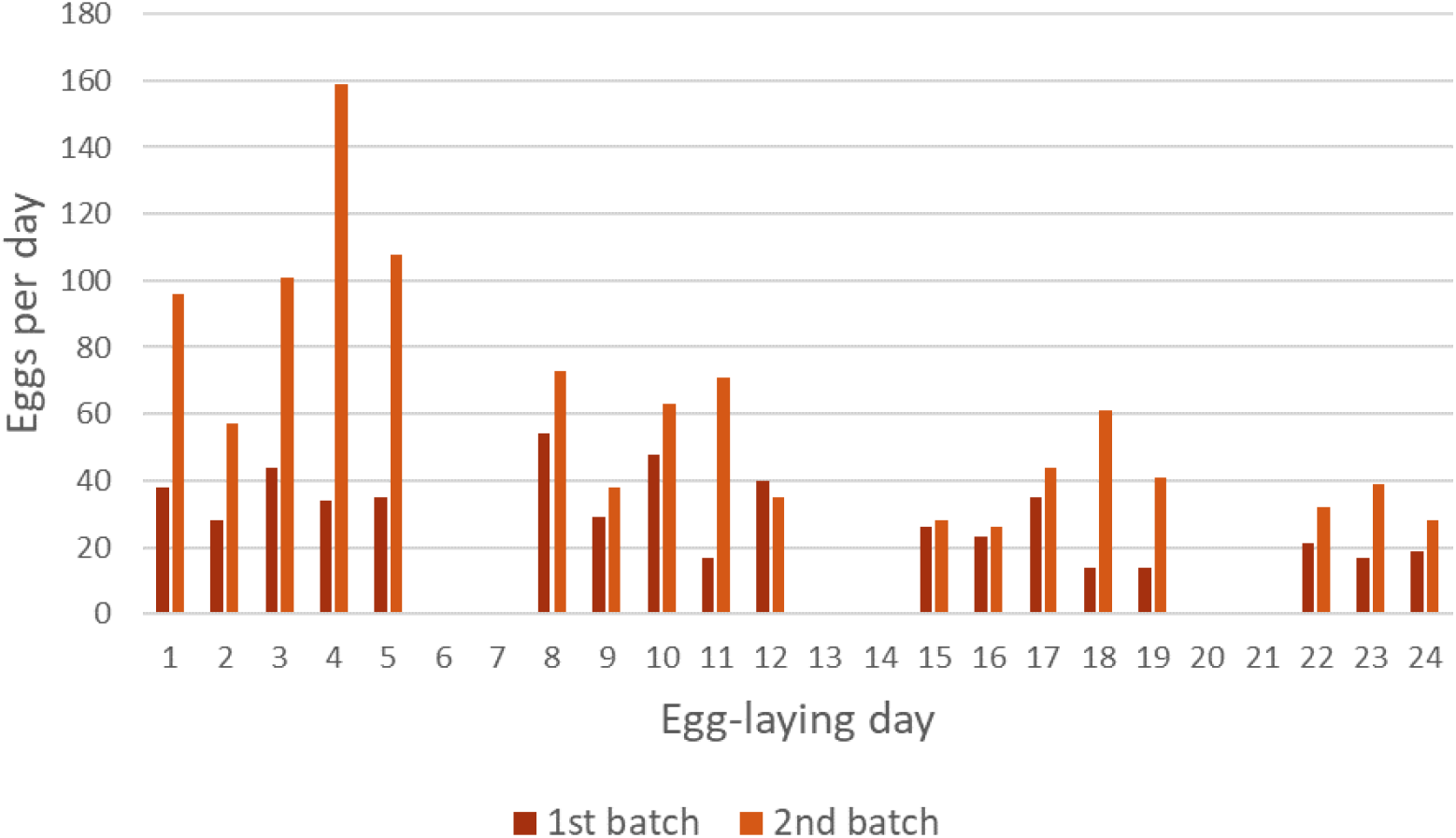
Daily egg yield decreased during the whole experiment. The number of eggs increased during the first week and then started to decline. The first and second egg-collection batches are shown with different colors. The first batch yielded 536 eggs and the second batch 1100 eggs.

**Figure 4:**
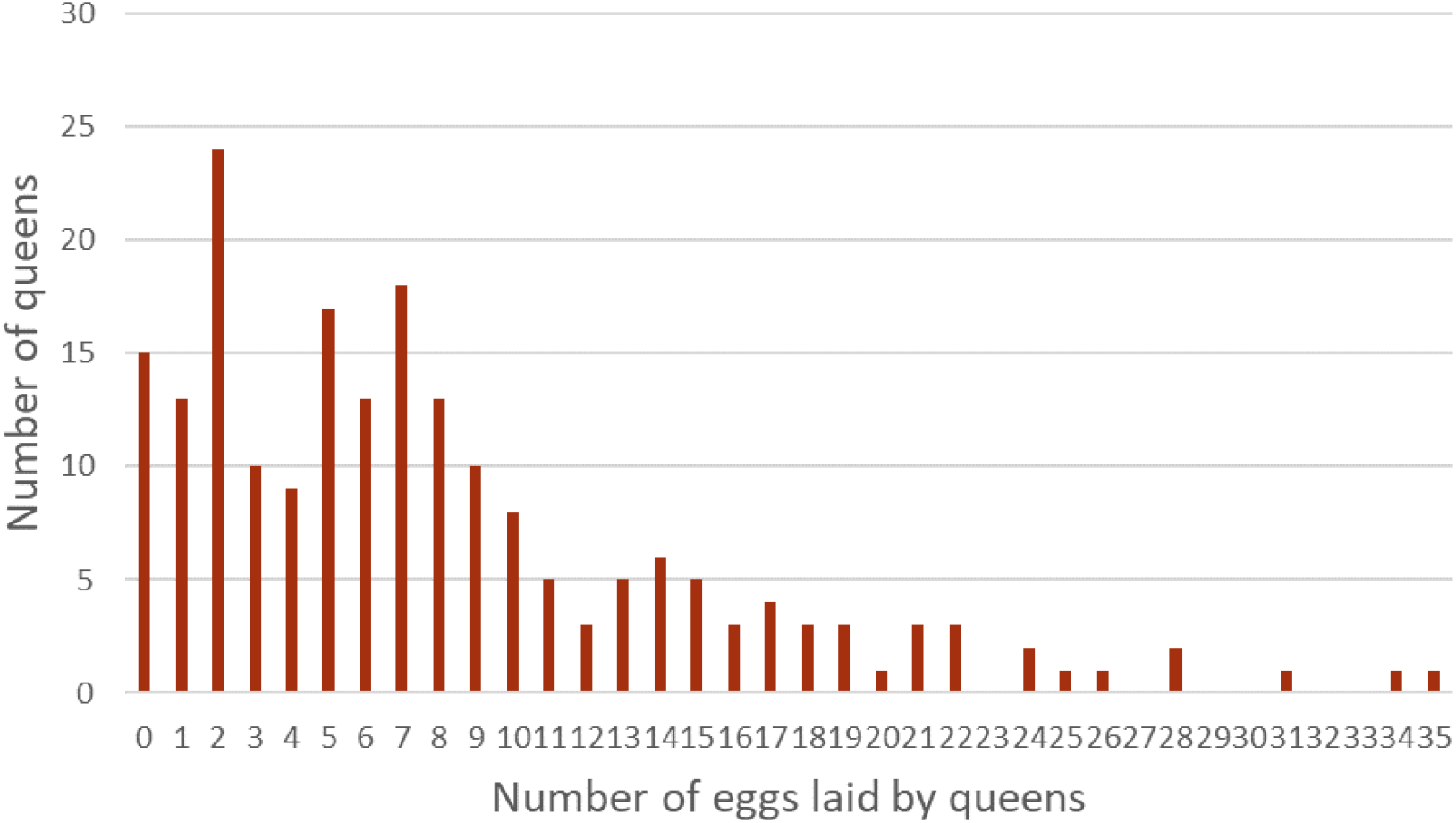
A large variation in egg yield per queen was observed. Fifteen queens did not lay any eggs and one queen laid 35 eggs.

### Concentration of CRISPR-Cas9 construct had a significant effect on the hatching success of eggs

There were in total 1624 eggs collected of which 1228 were injected with one of the three CRISPR constructs (R1, R2, R3), 195 were injected with sterile water and 201 were left untreated (Table 1). Mortality was high at the egg-stage where 85% of eggs did not hatch into larvae. Mortality at the larval stage was difficult to determine accurately because we could not determine if one larva had died and been replaced with a newly hatched larva. In total, circa 246 larvae hatched resulting in a 15% hatching probability over all treatments.

**Table 1:** Summary of the number of eggs treated and the outcome of the treatments.

|  |  |  |  |  | Successfully sequenced |  | Expressed genotype |  |
| --- | --- | --- | --- | --- | --- | --- | --- | --- |
| Construct | Concentration ng/μl | Total number of eggs | Total number of larva | Total number of adult workers | Larva | Workers | Mutated | Wild type |
| R1 | 80 | 204 | ca. 33 | 1 | 1 | 1 | 2 | 0 |
|  | 160 | 205 | ca. 7 | 0 | 2 | 0 | 1 | 1 |
| R2 | 80 | 245 | ca. 50 | 3 | 1 | 2 | 0 | 3 |
|  | 160 | 217 | ca. 45 | 6 | 2 | 4 | 2 | 4 |
| R3 | 80 | 165 | ca. 27 | 1 | 2 | 0 | 1 | 1 |
|  | 160 | 192 | ca. 18 | 4 | 1 | 4 | 1 | 4 |
| Water | N/A | 195 | ca. 21 | 1 | 19 | 1 | 0 | 20 |
| Contr | N/A | 201 | ca. 45 | 0 | 11 | 1** | 0 | 12 |
| Total |  | 1624 | ca. 246 | 16 |  |  |  |  |

\*\*Separate untreated sample

Out of the 1228 CRISPR-treated eggs 180 larvae (15%) hatched and out of the 195 water-treated eggs 21 larvae (11%) hatched. Hatching rate did not differ significantly between these two treatments (Fisher’s exact test, P=0.2233, Table 2). Therefore, injection of CRISPR-Cas9 construct did not increase mortality compared to water injection. When comparing the two different injected CRISPR-Cas9 concentrations, 18% hatched from the 80 ng/µl treatment and 11% hatched from 160 ng/µl treatment, the difference being significant and thus, higher concentration increasing mortality (Fisher’s exact test, P=0.0065, Table 2). Injection itself had a significant effect on egg mortality when comparing uninjected (22% hatched) and water-injected eggs (11% hatched). Uninjected eggs were more likely to hatch into larvae (Fisher’s exact test, P=0.0110, Table 2).

**Table 2:** Effect of different treatments.

| Fisher's exact test |  |  |  |  |
| --- | --- | --- | --- | --- |
|  | Eggs | Larvae | P-value | Conclusion |
| CRISPR | 1228 | 180 | 0.2233 | CRISPR does not increase mortality compared to |
| Water | 195 | 21 |  |  |
| 80 ng/μl | 614 | 110 | 0.0065* | Higher CRISPR-Cas9 concentration increases |
| 160 ng/μl | 614 | 70 | | |
| Water | 195 | 21 | 0.011* | Injection increases mortality |
| Uninjected | 201 | 45 |  |  |

\*Statistically significant P-value ( $P<0.05$ ).

In total 16 individuals matured into adult workers. In the uninjected control no adults emerged, and in the water-injected control one adult emerged. Similarly, one adult emerged from the R1-construct treatment, nine in the R2-construct treatment and five in R3-construct treatment (Table 1).

### Three gene-modified individuals showed a different eye color compared to the wild type

Sixteen adult workers were photographed (Figure 5) of which 14 were CRISPR-Cas9 treated, one was water-injected control, and one was an untreated individual outside of the experiment because uninjected controls did not produce any adult workers. Three out of the 14 CRISPR-treated individuals showed abnormal eye color compared to the black or very dark brown wild type. Two individuals had both eyes abnormal, and one individual had one normal and one abnormal eye. Inactivation of the *cinnabar* gene was expected to cause bright red eyes but instead, the eyes of the mutant individuals could be described as translucent with a similar lighter brown color as the surrounding cuticle with the white core of the eye visible. From the 1228 CRISPR-treated eggs three (0.0024%) had a genetic modification that could be scored from the phenotype.

**Figure 5:**
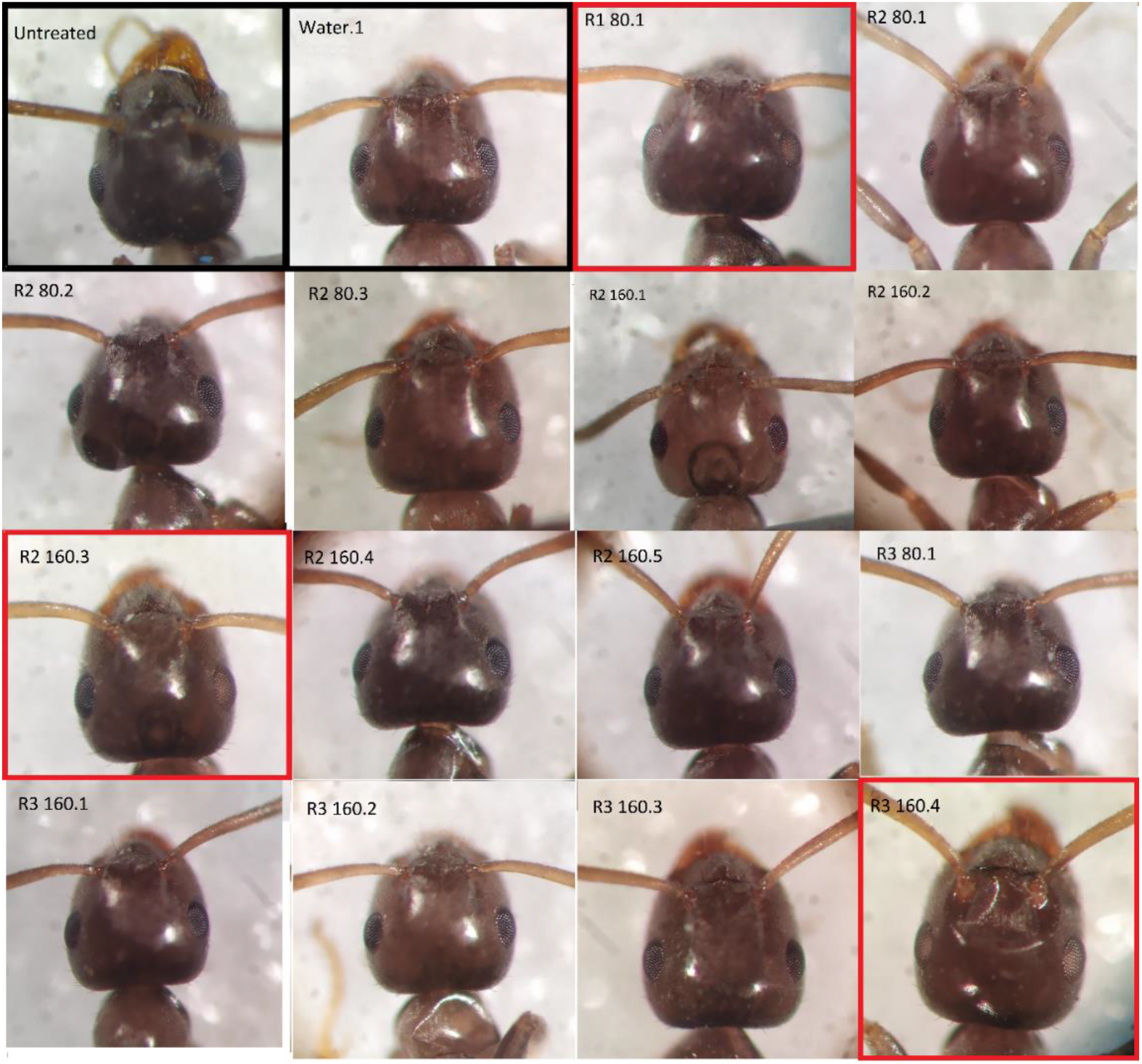
Eye-color phenotype of gene-modified and wild type ants. On the left of the top row is an untreated wild type individual and next to it a water injected control individual (black boxes). All the others are CRISPR-Cas9 treated individuals. Mutant individuals with visible phenotype are indicated with red boxes. R1 80.1 and R3 140.4 have both eyes abnormal and R2 160.3 has right eye abnormal while the left eye looks like a wild type eye. Wild type eyes are black or very dark brown and mutated eyes are whitish brown.

### Seven modified individuals carried mutations in the gene sequence

The target gene was sequenced from a total of 25 CRISPR-treated individuals (including 12 larvae and 13 adults) and 35 control individuals (GenBank accession numbers ON333547-ON333606). Two of the sequenced control individuals were adult workers (one water-injected and one uninjected) and 33 were larvae (20 water-injected and 13 uninjected). The obtained gene sequences were compared to *L. niger cinnabar* gene (GenBank accession: LBMM01007519.1). Thirty-two control individuals were successfully sequenced, none of which showed mutations. Seven mutated individuals were observed out of 20 successfully sequenced CRISPR-treated individuals (35%) (Table 1). Four individuals (one adult and three larvae) injected with R1-construct were successfully sequenced. Three of these four sequenced individuals contained 3-9 bp indels including the adult (R1 80.1) which also had abnormal eye color (Figure 5) and two larvae (R1 80 T2 and R1 160 T2). Nine successful sequences were obtained from individuals injected with R2-construct (6 adults and 3 larvae). Two of these individuals contained mutations: R2 160.3 (adult) carried a 6 bp deletion and also had abnormal eye color in one eye (Figure 5), and R2 160 T2 (larva) could not be sequenced due to mosaicism so the correct location of the mutation was confirmed by studying the chromatogram. Seven successful sequences were obtained from individuals injected with R3-construct (4 adults and 3 larvae). Two of these contained mutations: R3 160.4 (adult) carried a 4bp deletion and had abnormal eye color and R3 80 T1 (larva) was a mosaic carrying 4 and 12 bp indels.

In summary, seven out of the 20 sequenced CRISPR-treated individuals carried mutations compared to the 32 sequenced control individuals with no mutations (Fisher’s exact test, P=0.0026) indicating that the mutation rate between CRISPR-treated and untreated individuals differed significantly and that our protocol was successful in inducing mutations in *cinnabar*.

## DISCUSSION

In this study we successfully developed a protocol to rear and genetically modify *L. niger* ants with CRISPR-Cas9 technique. If gene-modified eggs are successfully reared to queens or males and these individuals reproduce, the mutations will become heritable. This can be utilized to breed ant strains for scientific research. Methods developed in this study can be applied to other ant species and other eusocial insects whose eggs and larvae do not survive without the care of the colony.

### Comparison to other CRISPR experiments conducted on ants and *Nasonia*

In terms of the life cycles of the ant species, the most suitable for comparison with *L. niger* is the imported fire ant *S. invicta* used by Chiu *et. al*., 2020, since both the species have morphologically distinct queen and worker castes. In contrast, *O. biroi* reproduces parthenogenetically and in *H. saltator* isolated workers become gamergate pseudo-queens able to reproduce. With these two species it is easy to create mutant strains while in *L. niger* CRISPR-treated eggs must first develop to queens or males to render the mutations heritable.

When compared to the four other CRISPR experiments conducted on ants (Trible *et. al*., 2017; Yan *et. al*., 2017; Chiu *et. al*., 2020; Sieber *et. al*., 2020), there are pros and cons in our approach. In all the four previous experiments the eggs were attached for injection using double-sided tape. The use of double-sided tape made the manipulation of the eggs very difficult after the injection which is why we settled for supporting the eggs against the petri dish wall. This facilitated the transfer of the eggs to the care of nursing workers, so the eggs did not need our monitoring until hatching. In Trible’s *et. al*. (2017) and Chiu’s *et. al*. (2020) experiments the eggs were left to develop on top of the tape until hatching after which Chiu *et. al*. (2020) sequenced the larvae because they could not rear them to adults. Trible *et. al*. (2017), on the other hand, managed to rear the larvae to adults using nursing workers and acclimating them using practice eggs just like in our experiment. In their experiment about 60% of the uninjected eggs hatched on petri dishes (our hatching rate was 22% with nursing workers) and about 50% of the larvae survived with adult nursing workers (none of our uninjected larvae matured to adulthood, while one water injected (0,5% of the eggs) and 15 CRISPR-injected (1,2% of the eggs) matured). Yan *et. al*. (2017) and Sieber *et. al*. (2020) removed the eggs from the tape after injection, washed them with ethanol and incubated them on agar plate treated with antibiotics to prevent bacterial or fungal growth. These studies also found out that the workers will eat the injected eggs and only after the first larvae had hatched the remaining eggs and larvae were transferred to a nest box with a few young workers. These studies did not mention anything about using practice eggs to train the workers which can explain the high mortality of eggs. Even though our methods of injecting and rearing the eggs were simpler and easier to carry out, we could not accurately determine the hatching rate because the nursing workers disposed of bad quality eggs and moved viable eggs around to make accurate counting difficult.

Our research methods were largely based on the work by Li *et. al*. (2017) where they modified the *cinnabar* gene of the parasitic wasp *N. vitripennis* with CRISPR-Cas9. Following their protocol, we injected the eggs with CRISPR-Cas9 concentrations of 80 ng/µl and 160 ng/µl, because they found these concentrations to be the most optimal in minimizing mortality and maximizing mutation rate. With smaller concentrations individuals had higher survival rate but the proportion of mutants was lower. With higher concentrations mortality was higher but those who survived had a higher proportion of mutants. In *Nasonia* 46% and 38% of the injected eggs survived to adulthood with 80 ng/µl and 160 ng/µl injection concentrations respectively, while in *Lasius* 0.8% and 1.6% eggs survived to adulthood with the same concentrations. In our study an unidentified factor hindered the larvae from maturing to adults, so it would be more meaningful to compare the survival rate in *Nasonia* to our hatching rate of eggs which was 18% and 11% with the same concentrations. Here it is noteworthy that *Nasonia* and *Lasius* have quite different life cycles: *Nasonia* larvae develop on their own inside host fly pupa while *Lasius* larvae are dependent on adult workers.

The eye color of the mutated ants caused some confusion. In Li *et. al*. (2017) the mutated *Nasonia* eyes were bright red like one could expect based on *Drosophila* research (Beadle & Ephrussi, 1937). When the *cinnabar* gene is inactivated, the production of dark brown ommochromes is blocked and only red pteridines are left to color the eyes. While the wild type eyes of *L. niger* ants are black or very dark brown, the eyes of the mutated ants were light brown and translucent. The color was similar to the surrounding cuticle and under the eye surface the white core was visible. It is possible that *L. niger* does not produce pteridines in the same scale as *N. vitripennis* or the pteridines produced by *L. niger* are different in structure and therefore more yellowish in color.

### Future outlook

Our protocol could be further optimized. The fifth hour produced the highest number of eggs, more than the four previous hours combined. In the future, eggs could be collected in six or seven-hour periods and study the effect of the extended egg-laying period on the egg yield.

The declining egg numbers during the whole experimental period could be caused by two circumstances. First, in Trible *et. al*. (2017) the egg-laying of the ants was synchronized by first letting the queens lay eggs in the nest and then removing the eggs for the experiment after which the queens started to lay new eggs again, i.e. the egg-laying occurred in the nest from where eggs were constantly removed. In our study eggs started to accumulate in the colony nests which might have resulted in the declining egg yield in the experimental egg-laying dishes when the experiment proceeded. If the extra eggs would have been removed from the colony nest, the egg-laying rate could have been maintained at high level during the experiment, if the queen regulates her egg-laying based on the amount of eggs present in the nest. Second, queen activity could have been affected by light-dark cycle in the climate chamber. We used 14-10 light-dark cycle but after discussing with a colleague we found out that in Finland insects thrive best in much longer light periods. Therefore, the light cycle could be optimized in future studies.

During the experiment we got in total only 16 adult workers from ca. 250 larvae hatched, which is a surprisingly low number regarding the total time used (93 days). At first, larvae pupated as expected but towards the end of the experiment the larvae growth rate and pupation seemed to slow down. The rearing nests were observed for 93 days after the egg collection ended which is almost twice as long as the average development time, and during the last 25 days no new pupa emerged. The low emergence of adults may be due to the larvae entering a diapause. According to Kipyatkov *et. al*. (2004), diapause occurs in northern populations more easily and to a larger extent when temperatures are lower.

In the future the growth rate and pupation of larvae could be studied in higher temperatures. In our study the eggs were reared in 23°C, while Trible *et. al*. (2017) and Chiu *et. al*. (2020) incubated their eggs in 25-30°C. Higher temperatures are known to speed up *L. niger’s* development time (Kipyatkov *et. al*., 2004) but we ended up using 23°C, because in 27°C the plaster bottom of the nest and drinking water cups dried much faster increasing the risk of eggs and workers dying. This could be avoided by using airtight containers as nests instead of petri dishes or by using thicker plaster layer and more water cups to retain moisture.

The effect of temperature on colony establishment period could be studied as well. In our study 56% of the queens managed to establish a colony in 23°C. In higher temperatures workers could develop faster and the colony establishment rate could be higher. According to Kipyatkov *et. al*. (2004), in southern populations temperature has a smaller effect on egg development rates than in northern populations. Due to their adaptation to shorter summers, northern populations start developing in lower temperatures and grow more intensively in higher temperatures compared to southern populations. Kipyatkov *et. al*. (2004) collected samples from St. Petersburg and Borisovka regions which are considerably more south than Oulu. In higher temperatures samples from Oulu could grow even faster than those from St. Petersburg. It can be also considered if our 2-month hibernation period before the experiment was sufficient for the ants. For example, in their overwintering experiment Haatanen, Ooik & Sorvari (2015) kept their ants in winter temperatures for over six months. In general, winter in Oulu lasts about that long. Our ants started to lay eggs after the 2-month hibernation period which was our aim but the effect of hibernation time on the egg-laying duration throughout the experiment could be studied further.

In summary, we successfully developed a protocol of genetic modification with CRISPR-Cas9 for the ant *L. niger* using easily detectable and non-lethal gene *cinnabar*. Our method can now be utilized to conduct more challenging experiments on *L. niger* targeting more vital genes for the ants’ viability or several genes simultaneously. In addition, this protocol can be referred to when developing gene editing methods for other ant and eusocial species.

## EXPERIMENTAL PROCEDURE

### Collecting the ants from the wild

1. To establish colonies, we collected young *L. niger* queens after their nuptial flight in 20^th^ and 21^st^ of July 2020. Mated queens are easy to recognize because they have dropped their wings. Queens with wings should not be collected.
2. The queens were collected from the surrounding areas of University of Oulu. *Lasius niger* ants are plentiful on roadsides and cracks of asphalt for example on parking lots and sidewalks and seams of courtyard tiling. Nests are easy to identify from small sand mounds around the cracks and seams of paved areas. We collected mated queens into small 5 ml plastic test tubes (anthouse.es) and sealed them with 1,5 ml plastic tubes filled with water and closed with cotton balls so the queens could drink water through the moist cotton. In total we collected 286 mated queens.
3. In year 2019 nuptial flight occurred in 20^th^ of July during which we collected 150 queens. In January 2021 there were left 66 of these colonies.

### Colony establishment in laboratory

1. We placed the queen tubes into a plastic box and put the box in a climate chamber (Sanyo) in 23°C, where the queens started to lay eggs.
2. The queen tubes were checked once every two weeks. During the check the tubes were opened, and the dried cottons were pushed deeper into the water tubes to ensure water supply. At the same time, we changed moldy tubes to clean ones and monitored the development of larvae.
3. The first larvae emerged in about two weeks and the first pupa about four weeks after the collection. Dead queens were removed from the box.
4. When the first workers emerged from their pupa and started to walk, we emptied the tubes to a 9 cm petri dish, hereafter called a “colony nest”, and started feeding them three times per week. Before this, the queens do not eat anything, and survive with their energy reserves collected before the nuptial flight.

### Queen colony maintenance

1. A colony was established for every queen, in which they lived during the experiment and where they could continue laying eggs with the help of workers. Colony nests were established in 9 cm petri dishes with ca. 5 mm layer of plaster cast on the bottom.
2. During the casting process a 1.5 ml plastic tube or a block of Lego was set in the wet plaster to create a depression in which the ants could gather. The plaster was moistened with water and an Eppendorf tube cap was placed in the nest to serve as a water cup. Food was dispensed on a 2×2cm aluminum foil piece and placed on top of the depression in plaster to create a dark cave for ants to hide.
3. The ants were fed three times a week and kept them at 23°C during the whole experiment. The carbohydrate-containing food was given in ca. 5mm lumps, and the mealworms were chopped into halves/quarters (see below the description of food). The water cups were filled, and the plaster moistened. We monitored the plaster moisture by observing the moisture of the food. If the food was dried up, the plaster was moistened more, and if the food was wet or molded, the amount of water was decreased. In general, 1-5 drops of water should be added to the plaster depending on the moisture levels. In case of mold, we cleaned the nest by wiping the plaster and dish cover with water and paper. If the mold was very abundant, the whole nest dish was replaced with a clean one. The queen, workers and eggs were carefully transferred to a clean nest using tweezers and a fine paint brush.

### Food

1. Three times a week we gave two types of food, that served as a source of carbohydrates and proteins. Carbohydrate-rich food was the Bhatkar diet (Bhatkar ja Whitcomb, 1970), which we prepared according to the recipe except without the vitamin-mineral-tablet. The food was cast on petri dishes and after solidifying, stored in a freezer for later use.
2. Mealworms (*Tenebrio molitor)* were fed to the ants to provide proteins. The worms were kept in a freezer, so they were easy to cut in pieces by hand when frozen.

### Hibernation

1. The queens’ egg-laying rate declined quite fast. Only a month after the colonies were established, we got only a few individual eggs from a preliminary experiment. We concluded that the colonies were preparing for winter, so they were transferred to a 4°C cold-room with the idea that after hibernation the queens would start active egg-laying again.
2. To prepare the colonies for hibernation, we removed food from the colony nests, and packed the nests into plastic boxes that were lined with moist Wettex-cloth. The box lids were not closed tightly to allow air circulation.
3. The colonies were checked once a week; the water cups were filled, moldy nests were changed to clean ones, and the Wettex-cloth checked for adequate moistness. The condensate water was removed from box lids and colony nests. Dead queens were removed. Colonies survive in cold-room conditions for about 3 months before mortality begins to rise. After that time period we found about 5-7 dead queens per week.
4. Before the experiment, we kept the colonies in the cold-room for two months, so the queens would be ready for a new “summer” and egg-laying season.

### Preparing the rearing nests

1. To conduct the experiment, we established eight rearing nests: six for different CRISPR-treatments and two for control-treatments. Three regions of the *cinnabar* gene were selected, and CRISPR-constructs (single guide RNA, sgRNA) were designed (see below ‘GuideRNA: designing, ordering and preparing**’**). Every construct was diluted in two different concentrations. Two controls were included: in the first control the eggs were injected with RNase-free water and in the second control the eggs were left untreated.
2. We gradually collected 20 workers into rearing nests when the colonies were transferred from the collection tubes to colony nests. We collected workers also from older colonies established in 2019 that had plenty of workers. Workers from different colonies could be mixed (we did not observe workers killing each other).
3. We collected a bunch of practice eggs and larvae from the colony nest and placed them in a rearing nest so the workers would assume the role of a nursing worker. Right before the experiment started, the practice eggs were removed from the nests so they would not be mixed with the CRISPR-treated eggs or controls.
4. The rearing nests were fed three times a week. Moisture levels were observed very carefully because eggs and workers cannot handle drought very well. In 23°C moisture levels could be maintained for about four days, but in the pilot experiment (not published) we observed that in 27°C the nests dried in 2-3 days. We prevented excess drought by adding more water cups in the nests. During the experiment, we kept the nests in 23°C and during the egg collection, the queens were in 27°C.

### Conducting the experiment

#### GuideRNA: designing, ordering and preparing

1. We determined *L. niger’s cinnabar* gene (GenBank accession: LBMM01007519.1) exon regions with the program GeneWise (Birney, Clamp & Durbin, 2004) by comparing the *L. niger* gene sequence to *Nasonia vitripennis* ortholog’s amino acid sequence (GenBank accession: XP_001602258.1). We identified nine exons in *L. niger* (Figure 6). The intron-exon structure differs between *L. niger* and *N. vitripennis. L. niger* exons have the following ranges: 4161…4230, 4548…4738, 4834…4882, 5643…5730, 5799…5970, 6069…6320, 6406…6495, 6699… 6839, 6930…7094.

**Figure 6:**
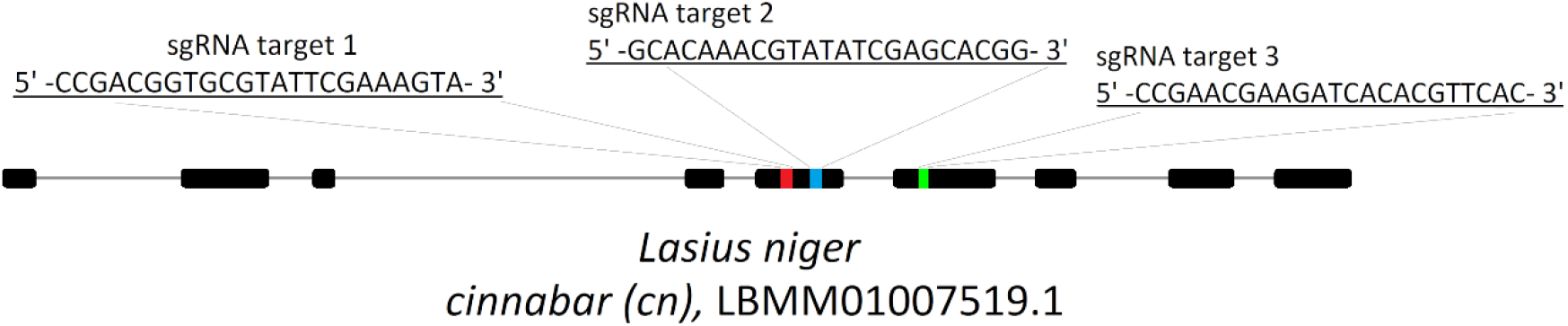
The relative size and location of exons and CRISPR-Cas9 targets in the gene *cinnabar*. Constructs R1 (red) and R2 (blue) are in exon 5 and construct R3 (green) is in exon 6.
2. We designed short guide RNAs (sgRNAs) to target three different parts of *L. niger* exons 5 (5799–5970) and 6 (6069–6320), in a hope that at least one of the sgRNAs would work effectively and disrupt the function of the *cinnabar* gene. SgRNAs were designed using RGEN Tools (rgenome.net).
3. We ordered TrueCut™ Cas9 Protein v2 and three different Invitrogen TrueGuide sgRNA Modified, custom sequences from Thermo Fisher Scientific. In this article those are referred to as R1, R2 and R3 constructs (Figure 6).
4. We diluted Cas9 protein and sgRNAs with RNase-free water to 160 ng/μl and 80 ng/μl and mixed them in equal proportions. These concentrations have been previously optimized for survival of the eggs and rate of successful mutations in the parasitic wasp *N. vitripennis* (Li *et. al*., 2017).

### Preparing the egg collection

1. We conducted the egg collection and the experiment in two batches due to the large number of colonies. The first batch was conducted in 11.1. - 3.2.2021 and the second batch in 8.2. - 3.3.2021. All the colonies were stored in a 4°C cold-room before egg collection. Colonies established in summer 2020 were placed in the cold-room on 30.10.2020 and colonies established in summer 2019 on 23.11.2020. For both batches we selected an equal amount of old (2019) and new (2020) colonies.
2. A week before the experiment, the colonies were moved from the cold-room to a 23°C climate chamber and fed three times a week, so that the queens had time to acclimate and start laying eggs.
3. We prepared containers for egg-laying. For each queen, we named a 6 cm petri dish with a number corresponding to their own colony. A semicircle shaped Wettex-cloth that covered half of the dish was placed on the bottom of the dish. The cloth was moistened with warm water so the conditions inside the dish would be favorable for the queen, and they had water to drink. The egg-collection dishes and water used for moistening the Wettex-cloths were stored in a 27°C climate chamber so they would always be in the correct temperature and the queens would not stress out for fluctuating temperatures.
4. Before collecting eggs for the experiment, the rearing nests were checked: were there enough workers and did they take care of the practice eggs. When needed, extra workers were added to the nests, in which case especially workers that actively cared for eggs were selected.

### Collecting the eggs for the experiment

1. Collection and injection of the eggs was carried out with a working pair. The queens were carefully picked with tweezers from the colony nest to their own egg-laying dishes. The egg-laying dishes were stacked on trays and transferred to a 27°C climate chamber. The dishes were checked once in an hour under a microscope. One of the working pair started to check the dishes and marked in a table how many eggs each queen had laid. If eggs were found, the queen was transferred to a clean dish to continue egg-laying in 27°C climate chamber. The other of the working pair started injecting the eggs. In this way the dish-checking and injecting could be done simultaneously.
2. We checked the dishes once in an hour to guarantee that the eggs were fresh, and no cell divisions would have occurred in the egg that would prevent the CRISPR-Cas9-construct from spreading evenly in all tissues. We continued the egg collection in the first batch for 4 hours and in the second batch for 5 hours, after which we returned the queens to their own colonies.
3. Egg collection was performed with the first batch for 3.5 weeks and after the egg-laying of the queens decreased to a minimum the second batch was taken out from hibernation and used for egg collection.

### Injection of the eggs

1. We injected the eggs under a microscope using PLI-100 Pico-Injector (Harvard Apparatus), mechanical micromanipulator and glass capillary needles (BioMedical Instruments ES-Blastocyst Pipette, spike, straight, inner diameter 12μm and length 55 mm). We thawed the sgRNA on ice and quickly spun it and loaded five microliters of the construct into the capillary needle. We injected the eggs with 10 PSI pressure for 80 milliseconds. Excess construct was kept in the tube on ice in case the needle needed refilling. After the day, excess construct was discarded.
2. Needle tip must penetrate the eggshell, so we supported the eggs against the petri dish wall so they could not roll around when injected. We moved the eggs around with a fine paintbrush moistened with water. This also helped to maintain proper moistness of the eggs. Manipulation of the egg was also easier with a wet brush.
3. After every injected egg, one drop of liquid was injected from the needle on the dish to make sure that the needle is not clogged. The needle used to inject the construct was discarded at the end of the day. The needle used for injecting the water control could be reused as long as it stayed functional. The needles were stored separately in their plastic containers.
4. During an injection day we injected only one construct and water. We injected the eggs collected during the day in following ratios: 60% of the eggs were injected with the construct, 20% with water and 20% were left uninjected. We changed the ratios midway of the experiment because control treatments started to accumulate too much in relation to CRISPR-treated eggs. The second half of the experiment was conducted with 80% construct injection, 10% water injection and 10% uninjected control. The needles and the construct were stored on ice when not used.
5. In one rearing nest there were always only one type of eggs (for example: eggs injected with construct R1 160 ng/μl had its own nest, water-injected their own etc.). Before the start of the experiment the practice eggs were removed from the rearing nests, so they were not mixed with treated eggs. The workers took care of the eggs by moving them in the right moisture conditions and removing damaged or unviable eggs. They also started to feed the larvae once they hatched and helped maturing workers out of their pupa. The rearing nests were kept in a 23°C climate chamber and fed like normal colonies three times a week. For every construct and control treatment we aimed to collect at least 100 eggs. Collection and injection of the eggs is depicted in Figure 7.

**Figure 7:**
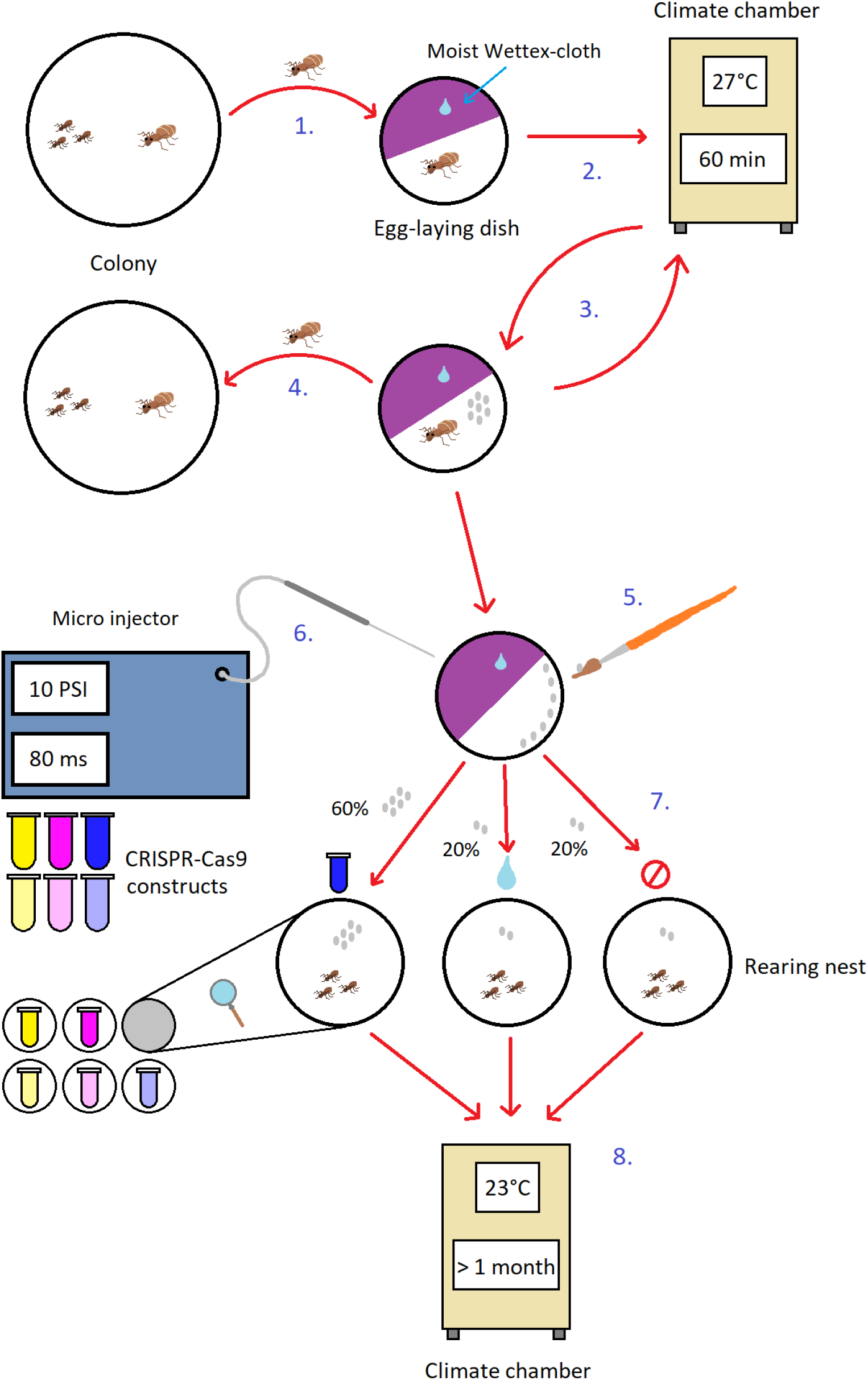
Schematic of the experiment. 1. The queen is picked from the colony to an egg-laying dish. 2. Egg-laying dishes are placed in a climate chamber for one hour. 3. After one hour the dishes are checked for eggs. If the queen has laid eggs, the queen is transferred to a new clean dish to continue egg-laying. 4. Egg collection is continued for 4-5 hours after which the queens are returned to their own colonies. 5. The eggs are prepared for injection by arranging them to a line against the dish wall using a fine brush. 6. A glass capillary needle is loaded with one of the six CRISPR-Cas9 constructs. 7. 60% of the eggs are injected with CRISPR-Cas9 construct, 20% are injected with water and 20% are left as an uninjected control. 8. The eggs are transferred to their own rearing nests according to their treatment so the workers can take care of them and rear them to adulthood.

### Monitoring the rearing nests

1. After the injection, the development of the eggs was monitored very closely. The day following the injection, we checked the nests to see if the workers had taken the eggs to their care. If the eggs had not been touched or the workers were clearly not interested in them, we moved a few workers with tweezers or paint brush directly on the eggs so they would be noticed. If this did not work, we transferred a few new workers from colony nests to rearing nests. For this, we selected large colonies and transferred only those workers that actively took care of the eggs to rearing nests. This usually prompted the rearing nest workers to take care of the eggs as well. The eggs were usually collected to large mounds and gathered in the same location where the workers spend their time. We counted the eggs regularly twice a week so we would notice if there were notable mortality of the eggs or larvae. During the count, we did not move the eggs to avoid damaging them. Therefore, the counting was not completely accurate, and the numbers were just approximates to estimate the mortality.
2. When pupae started to emerge, we checked the nests daily. There seemed to be two types of pupae of which the first one hatched in very early stage by assistance of adult workers, so the worker was not yet fully matured but was completely white and immobile. The worker then developed some pigment in its eyes, then mandibles and finally the rest of the shell before it started to walk on its own. The second type spent a longer time inside the pupa and emerged independently, almost fully matured and was ready to walk at the time of emergence. It still had a slightly lighter colored shell, so it was easy to identify from older workers.
3. At this point we transferred the new workers to a new nest to mature and put two older workers to take care of them. The older workers were marked with color paint on their abdomen so they could be separated from the CRISPR-treated individuals. We did the marking with paint markers by pumping little ink on a glass plate. Then we grabbed the worker by leg with tweezers and dipped a ball point pin into the ink and marked the ant carefully. This operation was easy to conduct in cold-room conditions, so the ants were much slower and easier to manipulate. The treated individuals were left to mature for 15 days, after which they were collected into individually marked Eppendorf tubes and frozen in –20°C. A few workers died before they reached the age of 15 days, so they were frozen at the time of death. After freezing, the workers were thawed and photographed under a microscope to identify the color of the eyes.

### Observing genetic modifications: Photographing the phenotype

1. To photograph the workers, we used the same microscope as during the injection (Olympus SZX-ILLD200). We thawed the individuals and placed them on a white Styrofoam block. The individuals were fastened to the foam by wedging them between two insect needles and correcting the head position with sharp tweezers so that both eyes were visible. The light source (Fiberoptic Heim L100) was pointed from both sides of the sample trying to minimize reflections from eyes. The samples were photographed with 50x magnification.

### Observing genetic modifications: Sequencing

1. We extracted DNA from workers (13 CRISPR-treated, 1 water-injected and 1 uninjected) and larvae (12 CRISPR-treated, 20 water-injected and 13 uninjected) selected for sequencing by using DNeasy Blood & Tissue Kit (QIAGEN) Purity and concentration of DNA samples were measured with a NanoDrop spectrophotometer (Thermo Fisher Scientific).

### Amplifying the target gene with PCR

1. We amplified the *cinnabar* gene using PCR and primers that were designed to cover different areas of the target gene. Primers were designed by using Primer3Plus (primer3plus.com). As a template for the primers, we used *Lasius niger cinnabar* gene (GenBank accession: LBMM01007519.1). SgRNA-constructs i.e., targets of the gene modification (red R1, blue R2 and green R3, Figure 6), were designed to target exons 5 and 6 (5799…5970 and 6069…6320), so the PCR primers were designed to cover these areas.
2. We designed in total five primer pairs (Table 3). Two pairs spanned a region from exon 4 to exon 7, one pair was designed closer to exons 5 and 6, one pair targeted a region in exon 5 and one pair in exon 6.

**Table 3:**
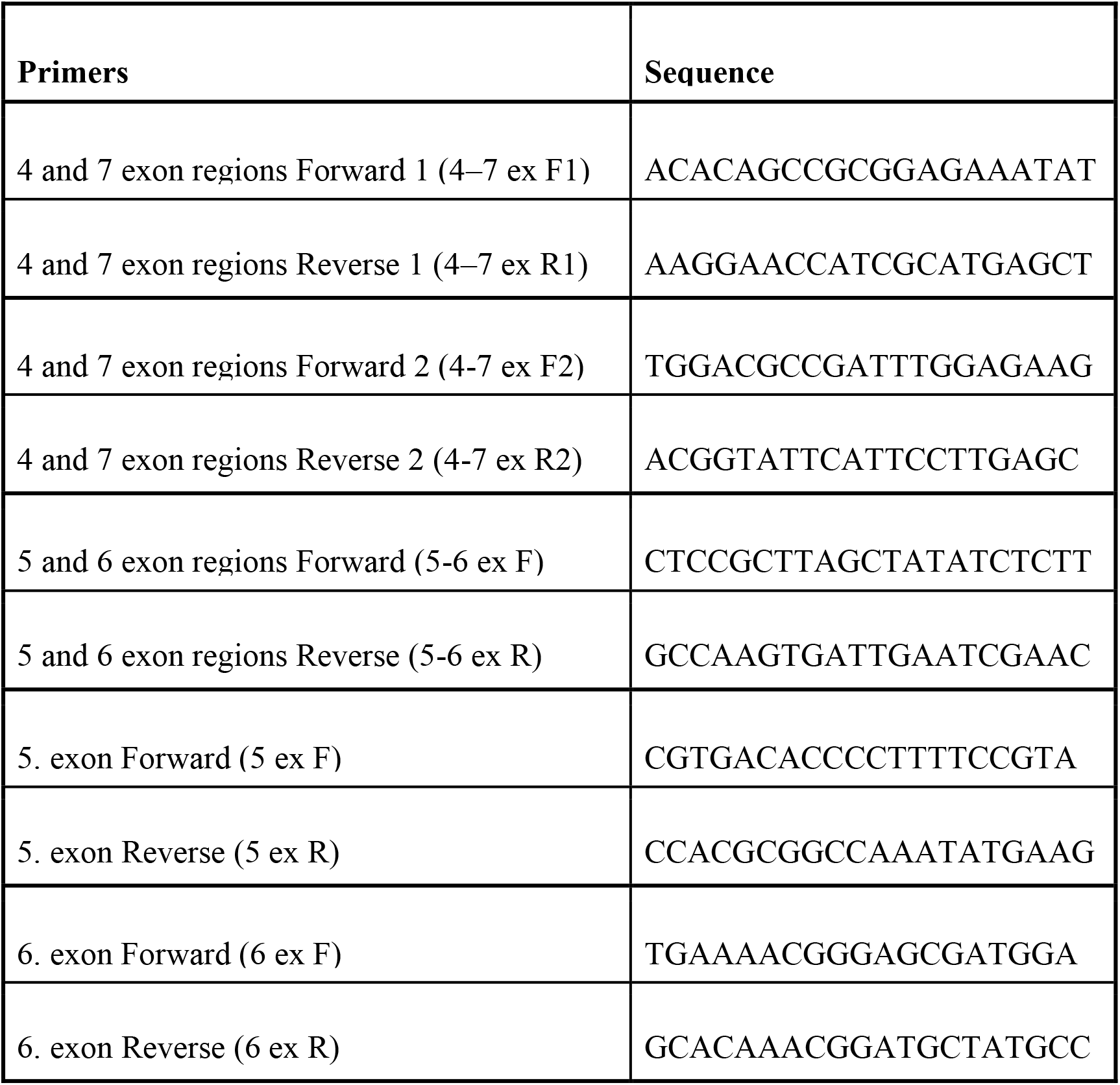
Primer pairs for PCR and sequencing.
3. PCR was conducted with following recipe:

4. The amount of template DNA and therefore the amount of water was changed according to DNA concentration so that the amount of DNA in PCR reaction would be 10-20 ng.
5. PCR parameters were the following:

3. PCR was conducted with following recipe:
|  |  |
| --- | --- |
| Dream Taq Green PCR Master Mix | 12,5 µl |
| F primer (25 µM) | 0,5 µl |
| R primer (25 µM) | 0,5 µl |
| Template DNA (n. 15 ng/µl) | 1 µl |
| Water | 10,5 µl |
| total | 25 µl |

6. We visualized the PCR results with agarose gel electrophoresis using 1.5% agarose gel and ran the samples with 100 V for 45 min. We tested all the PCR primers to be functional with unmodified samples. We conducted the PCR for the actual study samples with primers (4-7 ex F1) and (4-7 ex R1). Other primers we used for sequencing. For negative controls we used primers 5-6 ex F/R and water instead of DNA-sample to make sure that the reagents we used were not contaminated.

|  |  |  |
| --- | --- | --- |
| 95°C | 3 min |  |
| 95°C | 30 s |  |
| 55°C | 30 s | steps 2–4 were repeated 35 cycles |
| 72°C | 1 min |  |
| 72°C | 7 min |  |
| 4°C | ∞ |  |

### Sequencing the target gene

1. Purification of the PCR product from single-stranded DNA and dNTPs was carried out by Exol-FastAP-method:

2. We incubated the samples first in 37°C for 15 minutes. After this the enzymatic reactions were ceased by heating the samples to 85°C for 15 minutes. The samples were kept on ice before sequencing PCR.
3. For sequencing PCR, we selected the primers for sgRNA-constructs as follows:

|  |  |
| --- | --- |
| PCR-product | 5 μl |
| FastAP (1U/μl) | 1 μl |
| Exol (20 U/μl) | 0,5 μl |

- R1 and R2 constructs were sequenced with primers (4-7 ex F2) and (5 ex R)
- R3 construct was sequenced with primers (6 ex F) and (6 ex R)
- Water-injected and uninjected controls were sequenced with primers (4-7 ex F1) and (4-7 ex R1)
- The sequencing PCR was conducted with the following recipe:

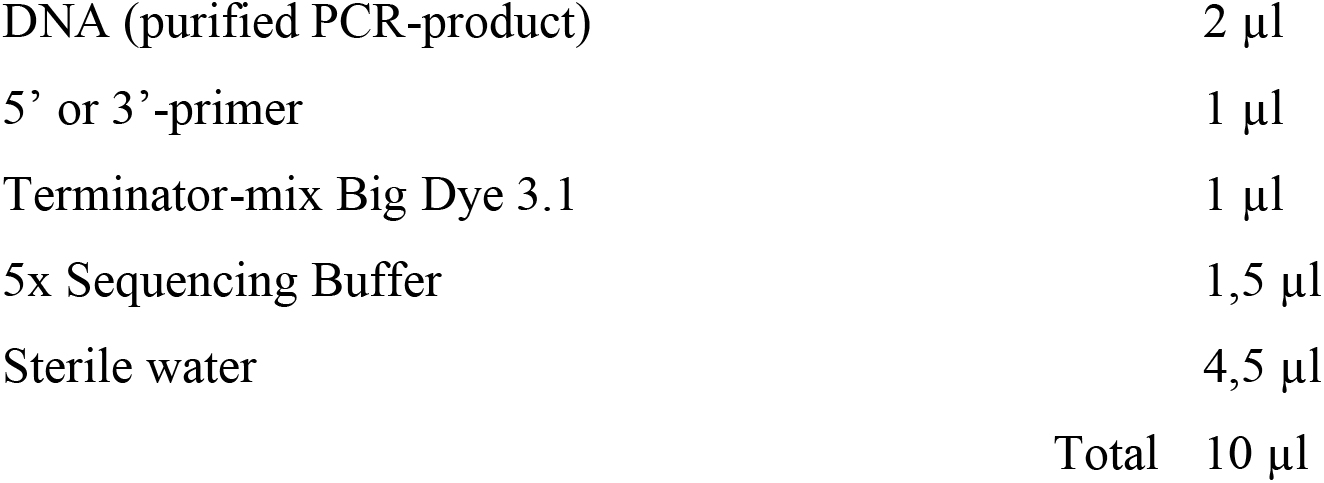
- The samples were pipetted to a PCR plate and sealed with a membrane. The plate was centrifuged and ran in PCR with the following parameters:

|  |  |
| --- | --- |
| DNA (purified PCR-product) | 2 μl |
| 5' or 3'-primer | 1 μl |
| Terminator-mix Big Dye 3.1 | 1 μl |
| 5x Sequencing Buffer | 1,5 μl |
| Sterile water | 4,5 μl |
| Total | 10 μl |

- Finally, the PCR products were purified from extra nucleotides by filtering it through Sephadex gel.

|  |  |  |
| --- | --- | --- |
| 96°C | 1 min |  |
| 96°C | 10 s |  |
| 50°C | 5 s | Stages 2–4 were repeated for 25 cycles |
| 60°C | 4 min |  |
| 4°C | ∞ | |

### Cloning

Four of our CRISPR-treated individuals contained mosaic mutations which rendered the sequence unreadable during the analysis. We separated the different alleles from each other by cloning them with TOPO TA Cloning Kit for Sequencing (Invitrogen) following the manufacturers protocol.

## ACKNOWLEDGEMENTS

We would like to thank Ville Salo for help with the injections. This study was funded by the Emil Aaltonen Foundation (LV), Academy of Finland grant number 343022 (LV), Societas Biologica Fennica Vanamo (MK) and the Kuopio Naturalists’ Society (MK). JK was funded by Academy of Finland grant number 328961.

